# Deep sequencing reveals two Jurkat subpopulations with distinct miRNA profiles during camptothecin-induced apoptosis

**DOI:** 10.1101/237099

**Authors:** İpek Erdoğan, Mehmet İlyas Coşacak, Ayten Nalbant Aldanmaz, Bünyamin Akgül

## Abstract

microRNAs (miRNAs) are small non-coding RNAs of about 19-25 nt that regulate gene expression post-transcriptionally under various cellular conditions, including apoptosis. The miRNAs involved in modulation of apoptotic events in T cells are partially known. However, heterogeneity associated with cell lines makes it difficult to interpret gene expression signatures especially in cancer-related cell lines. Treatment of Jurkat T cell leukemia cell line with the universal apoptotic drug, camptothecin, resulted in identification of two Jurkat sub-populations: one that is sensitive to camptothecin and the other being rather intrinsically resistant. We sorted apoptotic Jurkat cells from the non-apoptotic ones prior to profiling miRNAs through deep sequencing. Our data showed that a total of 184 miRNAs were dysregulated. Interestingly, apoptotic and non-apoptotic sub-populations exhibited a distinct miRNA expression profile. In particular, 6 miRNAs were inversely expressed in these two sub-populations. The pyrosequencing results were validated by real time qPCR. Altogether these results suggest that miRNAs modulate apoptotic events in T cells and that cellular heterogeneity requires careful interpretation of miRNA expression profiles obtained from drug-treated cell lines.

## Introduction

Apoptosis is programmed cell death triggered by various stimuli from outside or inside the cell such as ligation of cell surface receptors, treatment with cytotoxic drugs or irradiation and result in transcriptionally regulated activation of a number of regulatory proteins (Blank and Shiloh, 2007). Apoptosis is characterized by exposure of phosphotidylserine on plasma membrane outer leaflet, membrane blebbing, cellular shrinkage, chromatin condensation and fragmentation of nuclear DNA leading to formation of apoptotic bodies (Blagosklonny, 2000; Baumann et al., 2002).

T cells constitute a vital branch of cell-mediated immunity and homeostasis of the immune response is sustained through a balance between proliferation and apoptosis of T cells. A wealth of information has accumulated over the past two decades about the transcriptional regulation of genes mediating apoptosis in T cells. The recent discovery of small RNAs, however, suggests that post-transcriptional gene regulatory networks might have prominent effects on modulation of T cell apoptotic pathways (Lodish et al., 2008; O’Connell et al., 2010). miRNAs, one of those small RNAs, are non-coding small RNAs of about 19-25 nucleotides in length, which are transcribed by RNA polymerase II (Bartel, 2004; Bartel, 2009). Followed by nuclear processing by Drosha and cytoplasmic processing by Dicer, the mature miRNA strand negatively regulates gene expression by translational inhibition or destabilization of mRNAs (Miska 2005; Lawrie, 2007; Stefani and Slack, 2008). miR-14 and bantam were the first members of miRNAs shown to modulate apoptotic functions in *Drosophila* (Brennecke et al., 2003; Xu et al., 2003). Over the past few years, a clear link has been established between apoptosis and miRNAs (Su et al., 2015), particularly in cancer development (Kumar et al., 2007; Marcucci et al., 2011).

Besides the significance of miRNAs in T cell functions, miRNA-mediated T cell apoptosis has also been associated with OncomiRs (Oncogenic miRNAs) in leukemiagenesis (Calin et al., 2009; Pekarsky et al., 2009; Chen et al., 2010). In fact, miRNA profiling studies particularly on cancerous tissues have clearly shown that each leukemia type (for instance, CLL vs ALL) possesses a prominent miRNA expression signature (Zanette et al., 2007). Additional studies showed that these miRNAs regulate the expression of apoptotic or anti-apoptotic mRNAs (Mott et al., 2007; Xiao et al., 2008; Akao et al., 2009).

Although the use of cell lines has led to the identification of a number of dysregulated miRNAs involved in apoptosis and/or leukemiagenesis (Li et al., 2013; Yamada et al., 2014; Zhou et al., 2014; Fan et al., 2016), cellular heterogeneity associated with cancerous tissues require careful interpretation of the data acquired from human studies. Cellular heterogeneity is also an important issue that needs to be taken into account while interpreting the data collected from cell lines. It is well documented that cells use cellular heterogeneity to function properly and survive (Benchaouir, 2004; Stockholm et al., 2007; Walling et al., 2012). Jurkat cells were also reported to be heterogeneous as sublines from the same clones displayed different cellular morphologies and growth patterns (Snow and Judd, 2009). Although miRNA heterogeneity is known to exist across different cell lines that originate from the same cancer type (Lu et al., 2015), the potential for differential miRNA expression across the cells in the same cell line has not been reported before. To identify miRNAs that regulate apoptosis in Jurkat cells and also to test the effect of cellular heterogeneity on drug response and miRNA expression profiles, we triggered apoptosis in Jurkat cells with camptothecin, an inhibitor of DNA topoisomerase I and a potent inducer of apoptosis (Li and Liu, 2001; Pommier et al., 2003). Following the drug treatment, we sorted the apoptotic sub-population (as defined by Annexin V^+^) from the non-apoptotic sub-population by magnetic beads. Deep-sequencing of small RNAs isolated from each subpopulation revealed that each sub-population possessed a distinct miRNA expression signature that might be associated with population-specific apoptotic response.

## Methods

### Cell culture, drug treatment and transfection

Jurkat human leukemic T cells (American Type Culture Collection clone E6.1) were maintained in RPMI 1640 (Gibco) supplemented with 2 mM L-glutamine, 10% fetal bovine serum (GIBCO) and 100 U penicillin/100 μg streptomycin (Biochrom AG) in an atmosphere of 5% CO_2_ at 37^o^C. 10^6^ cells were treated with different concentrations of camptothecin (Sigma) and incubated at 37^o^C, 5% CO_2_ to determine the dose kinetics. Cells both in treatment and control group were labeled with PE-conjugated Annexin V and 7AAD (BD Pharmingen) and analyzed by flow cytometry (BD FacsArray) to identify stages of apoptosis. Cells that were Annexin V-positive yet 7AAD-negative were defined as early apoptotic ones. Annexin-V-positive cells were then seperated by Annexin V microbead kit (Miltenyi Biotech) according to the manufacturer’s instructions. Four fractions were obtained as follows: untreated Annexin-negative (JNN), untreated Annexin-positive (JNP), treated Annexin-positive (JAP) and treated Annexin-negative (JAN) cells. Because sufficient cells could not be obtained from the JNP cells, they were excluded from the study.

### Total RNA Isolation and Deep sequencing

Total RNA was isolated with mirVana miRNA isolation kit (Ambion) according to the manufacturer’s instructions. Total RNA samples were treated with TurboDNase DNA-free kit (Ambion) to remove traces of genomic DNA contamination. RNA integrity was determined by Bioanalyzer (Agilent 2100) using RNA 6000 Nano Kit (average RIN ~ 9-10).

3 replicates from untreated Annexin-negative (JNN), treated Annexin-positive (JAP) and treated Annexin-negative (JAN) cells were mixed in equal amounts and sequenced using Illumina Genome Analyzer by Fasteris (Switzerland). The fragments missing either adaptor were excluded from further analyses. The adaptor sequences of bp-15-29 inserts were removed prior to the subsequent data analyses. Nexalign program (Timo, Lassmann, unpublished data, http://genome.gsc.riken.jp/osc/english/software/src/nexalign) was used to align reads to rRNA (NCBI, U133169), hairpin and mature miRNAs (miRBase, R18) (Vaz et al., 2010). The sequences were aligned first for exact match and then the remaining sequences were used to identify matches with up to three mismatches as described previously (De Hoon et al., 2010). The data were deposited to GEO under the accession number GSE35442.

### Real-time qPCR Analyses

RNAs smaller than 200 nt were isolated directly from the cells using the miRVana miRNA isolation kit according to the manufacturer’s instruction (Ambion). cDNA was prepared from the small RNAs using RT^2^ miRNA cDNA kit (SA Biosciences). qPCR was performed in duplicates of three biological replicates (Roche, Lightcycler 480). U6 ncRNA was used for normalization.

## Results

### Identification of two Jurkat sub-populations with different apoptotic properties

We first performed dose-response (0.5-1024 μM) kinetics to determine the optimal drug treatment conditions for capturing cells at the apoptotic stage (Annexin V-positive and 7AAD-negative). When Jurkat cells were treated with camptothecin for 4h, apoptosis was observed in direct proportion to the dose applied up to 8 μM (Fig. 1A). Percentage of apoptotic cells did not significantly change and reached a plateau after 8 μM. Apoptosis was observed in 41% of the cells in camptothecin-treated group compared to 5.5% in the control cells (Fig 1A, *P* < 0.05).

**Figure 1.**
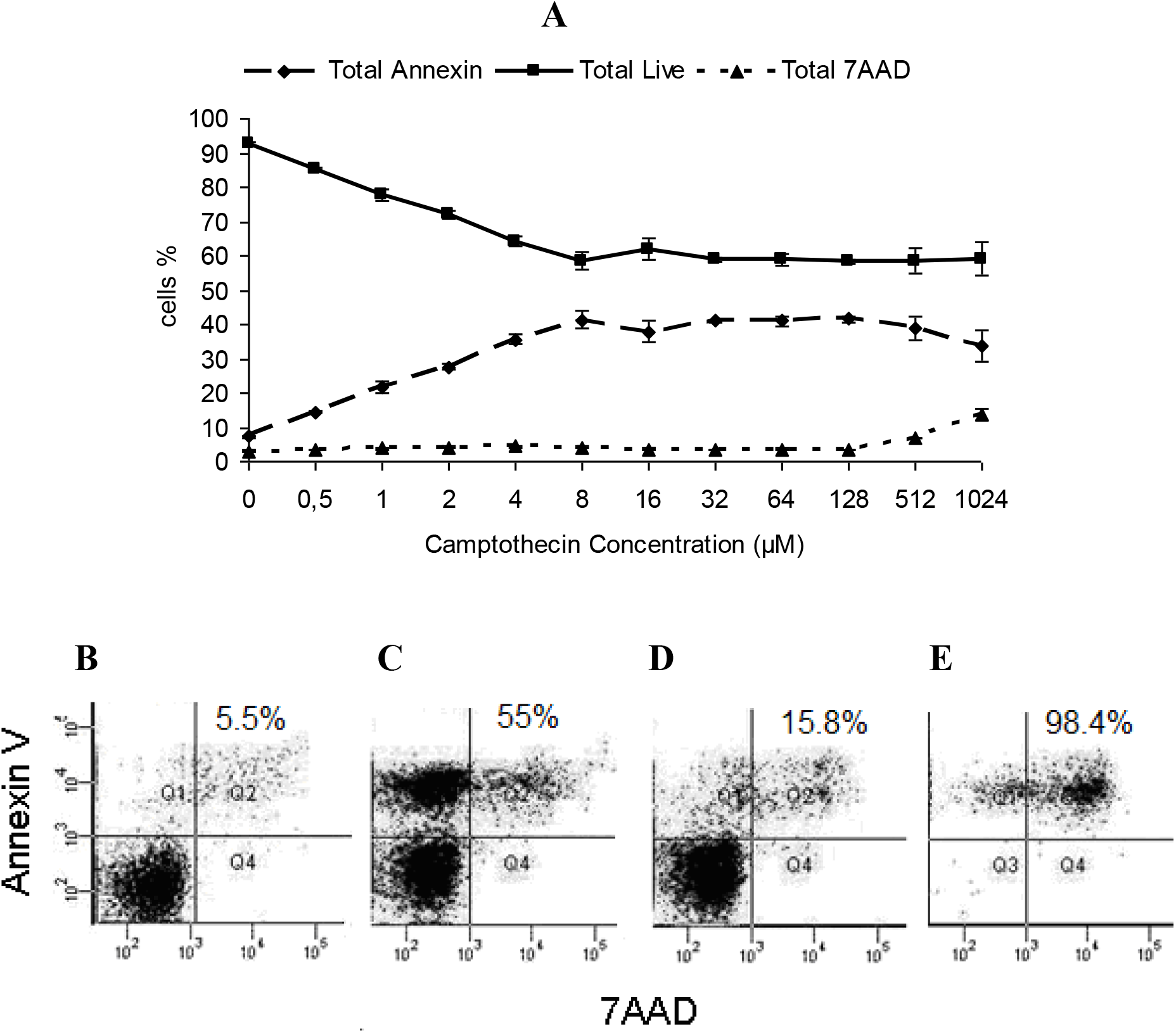
Dose response and identification of apoptotic cells by flow cytometry. A. Dose kinetics of camptothecin. Jurkat cells were treated with a range of camptothecin (0.5-1024 μM) for 4 hours and the apoptotic cells were determined with Annexin V/7AAD labeling. *(B-E)* Enrichment of apoptotic cells with magnetic bead separation. Jurkat cells were treated with 8 μM camptotecin for 4 hours and the apoptotic cells were sorted using an Annexin V magnetic bead separation kit. The apoptosis rate of the following samples was determined by flow cytometry: (B) control, untreated cells (JNN); (C) camptothecin-treated cells, (D) camptothecin-treated and magnetic bead-sorted Annexin V-negative cells (JAN); (E) camptothecin-treated, magnetic bead-sorted Annexin V-positive cells (JAP). It is important to note that the cells become Annexin V/7AAD-double positive following the sorting.

The unresponsiveness of some cells to the drug has led us to hypothesize that Jurkat cells may contain additional sub-populations, each of which might have a distinct camptothecin-mediated apoptotic property. Thus, we increased the concentration of the drug up to 1 mM (128-fold in excess of the maximum 8 μM concentration). Interestingly, over fifty percent of the camptothecin-treated cells still remained relatively resistant to the drug treatment despite over 100-fold camptothecin concentrations, suggesting the presence of a drug-resistant second subpopulation (Fig. 1A). The apparent difference in the apoptotic response could stem either from an uneven exposure of the cells to the drug or from intrinsic resistance of a sub-population to the drug. To ensure that the differential apoptotic capacity of two sub-populations is not due to the uneven drug treatment, we sorted the non-apoptotic cells and re-treated them with the drug. To this end, Jurkat cells were first treated with 8 μM camptothecin for 4h and the Annexin V-negative non-apoptotic cells (JAN) were separated from the apoptotic cells (JAP) using Annexin V-conjugated magnetic beads (Fig. 1B-E). Flow cytometry analysis of the sorted cells showed that the sorting efficiency was as high as 98.4% (Purity > 85 %, 95% in average) (Fig. 1E). Re-treatment of the non-apoptotic cells (JAN) with 8 μM camptothecin for 4h showed that this fraction of the cells was indeed intrinsically resistant to induction of apoptosis by camptothecin. 42.4% of the cells underwent apoptosis in the first round of treatment while the second camptothecin treatment triggered apoptosis only in 7.4% of the JAN cells (Fig. 2, *P* < 0.05). We concluded that there are intrinsic gene expression properties, e.g., miRNAs, associated with resistance to the camptotecin-induced apoptosis in Jurkat T cells apparently composed of at least two sub-populations.

**Figure 2.**
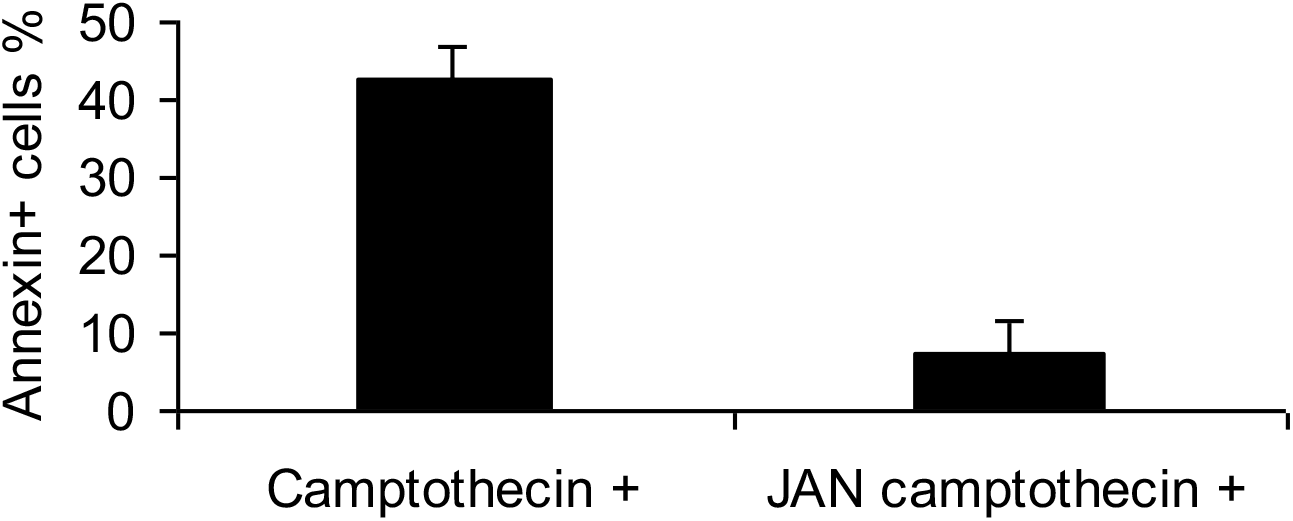
Two sub-populations of Jurkat cells with different camptothecin-mediated apoptotic capacity. Following the magnetic bead separation of camptothecin-treated and magnetic bead-sorted Annexin V-negative cells (Figure 1D, JAN cells), these cells were re-treated with camptothecin for 4 hours. Apoptosis was measured with Annexin V/7AAD labeling. Camptothecin +, cells treated with camptothecin once; JAN camptothecin +, sorted cells that were re-treated with camptothecin.

To investigate whether each Jurkat sub-population possesses a distinct miRNA expression profile, we compared the miRNA expression patterns in 3 replicates of each sub-population. The JNN sample contains the Annexin V-negative cells, which were not treated with the drug (negative control) whereas the JAN and JAP samples contain the Annexin V-negative (intrinsically camptothecin-resistant) and positive (intrinsically camptothecin-susceptible) cells, respectively, which were treated with the drug. It should be noted that although the majority of the camptothecin-treated cells were Annexin V-positive and 7AAD-negative prior to sorting, they shifted to a Annexin V/7AAD-double positive phenotype subsequent to sorting. We used the Illumina platform (Fasteris, Switzerland) to quantitatively measure the amounts of small RNAs in each sample. After the removal of the adapter sequences, 91.3% of all reads contained 15-29-bp inserts. Based on the size of the inserts, there appeared to be two major small RNA populations, one of 22-23 bp and the other 28 bp, each representing miRNAs and tRNA-derived small RNAs, respectively (Fig. 3). The alignment of the reads to the known RNAs revealed two striking points with respect to the small RNA content of each sample: (1) the control JNN cells are rich in miRNA, which constitutes 60% of small RNAs. The miRNA content plummets to 26 and 16% in the camptothecin-treated JAN and JAP samples, respectively; (2) the drug treatment induces major tRNA fragmentation, constituting up to 45% of all small RNAs (Fig. 3; nt-27-29 region). We did not notice a major difference in the proportion of other small RNA categories although there may be differences in the expression of individual small RNAs.

**Figure 2.**
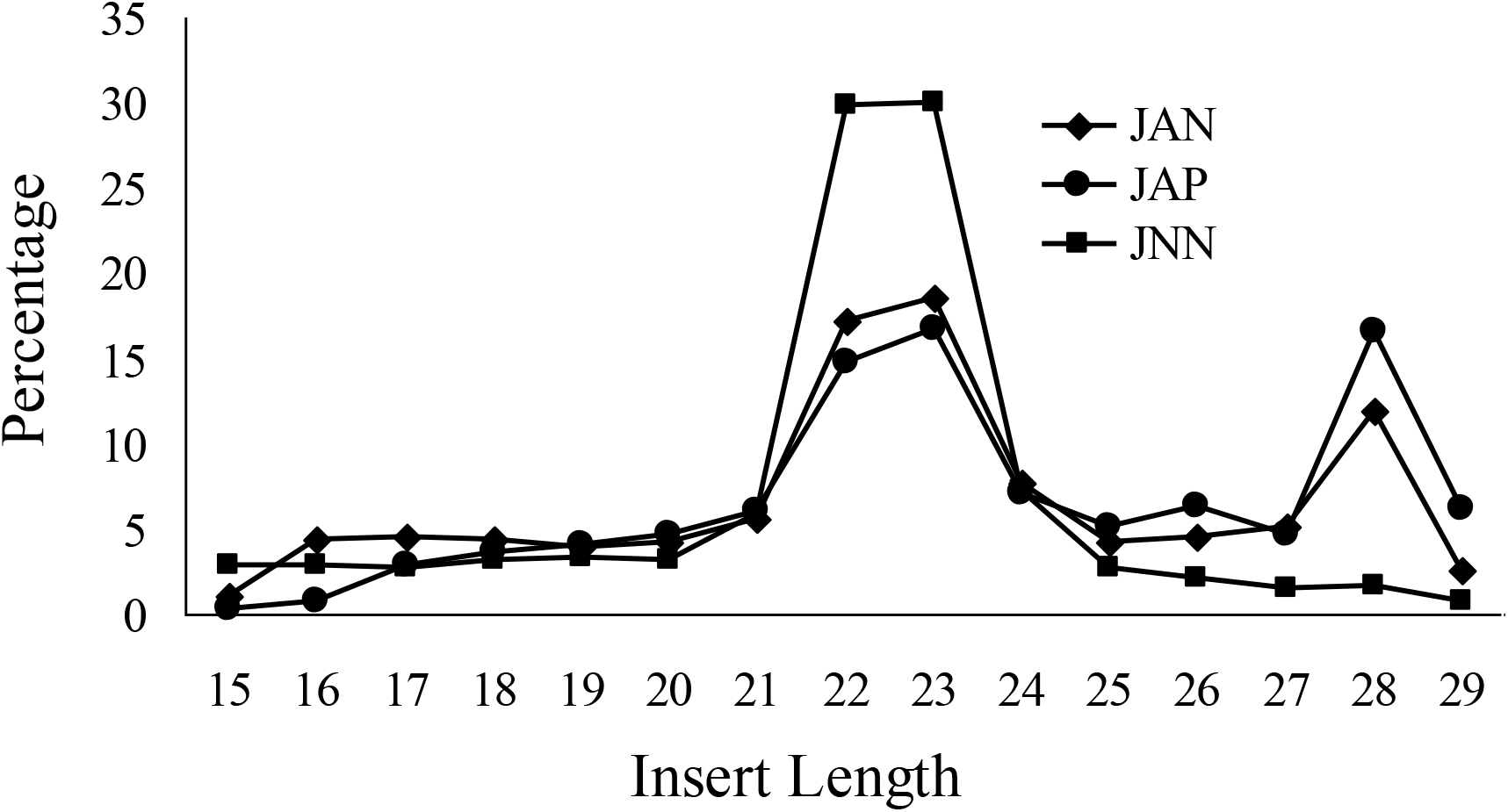
Small RNA profiles of camptothecin-treated Jurkat cells. A. The 15-29-bp fragments were sorted based on their size and plotted as percentage. JNN, small RNA population in the drug-free, control cells; JAP, camptothecin-treated and Annexin V-positive cells and JAN, camptothecin-treated and Annexin V-negative cells.

### Apoptosis is regulated by differential miRNA expression

The alignment of reads to the known miRNAs in miRBase (R18) resulted in identification of 184 miRNAs differentially expressed among the three samples (Table 1). Our list includes the differentially expressed miRNAs, whose expression are greater than 10 RPM (read per million) in all three samples. Thus, the number of differentially expressed genes could be greater. Camptothecin treatment usually suppresses miRNA expression compared to the control cells (Table 2A, 38 induced miRNAs versus 144 down-regulated miRNAs in the JAN or JAP samples). We identified a single miRNA, miR-1246, over-expressed in the camptothecin-treated JAN and JAP samples. However a total of 79 miRNAs were down-regulated in response to the camptothecin treatment. Interestingly, 16 and 30 miRNAs were down- and up-regulated in the drug resistant JAP sample whereas they were equally expressed in the drug sensitive JAP sample. More interestingly, a total of 6 miRNAs (let-7b-5p, miR-15a-5p, 324-5p, 128, 425 and 720) were reciprocally expressed in the drug-sensitive and resistant cells. To validate our findings from deep-sequencing, we randomly chose 7 miRNAs for validation by real-time qPCR, namely miR-7, 17, 18a, 25, 26a, 93 and 425. As shown in Table 2B, the qPCR results were quite consistent with the deep-sequencing data.

**Table 1.**
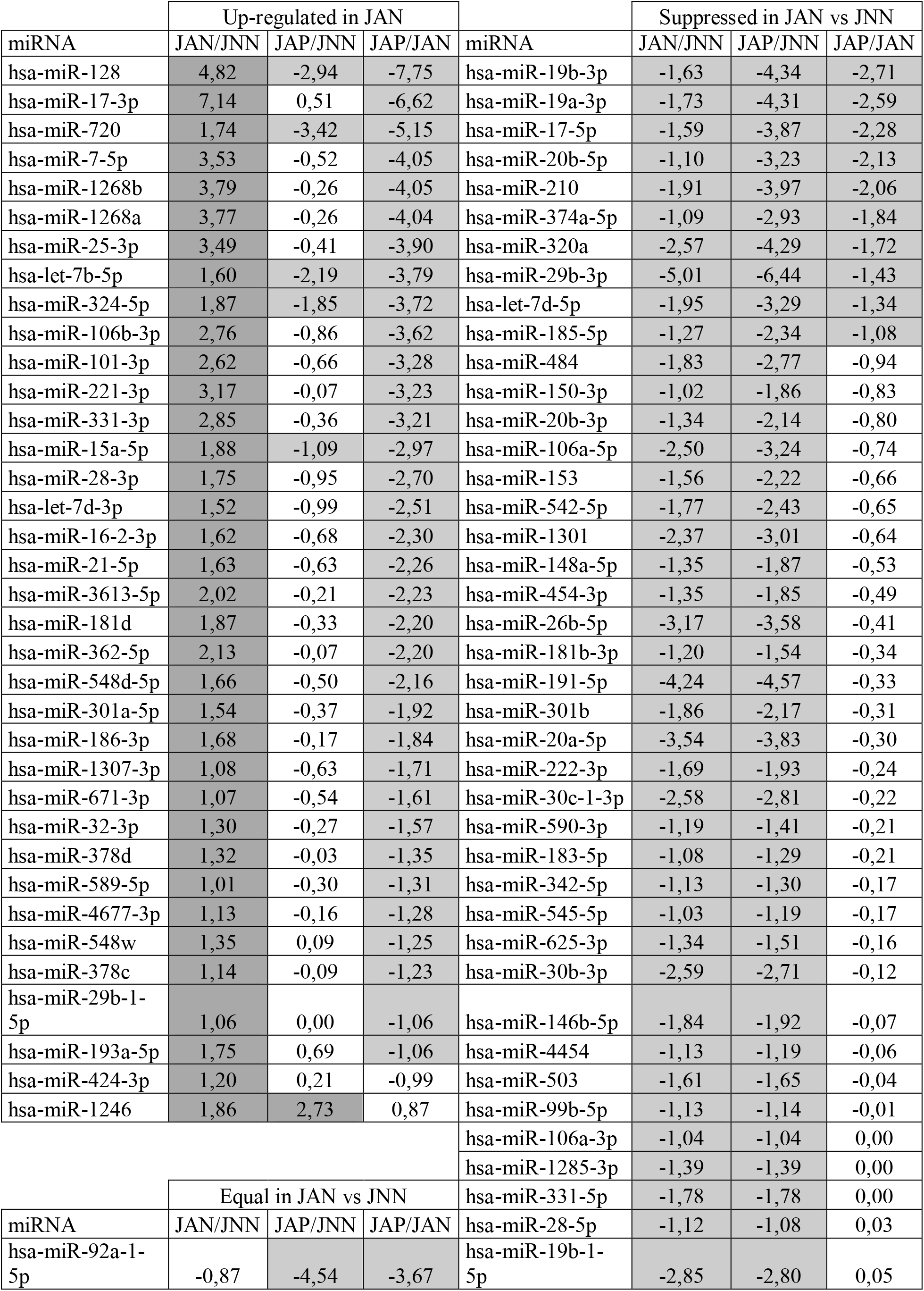
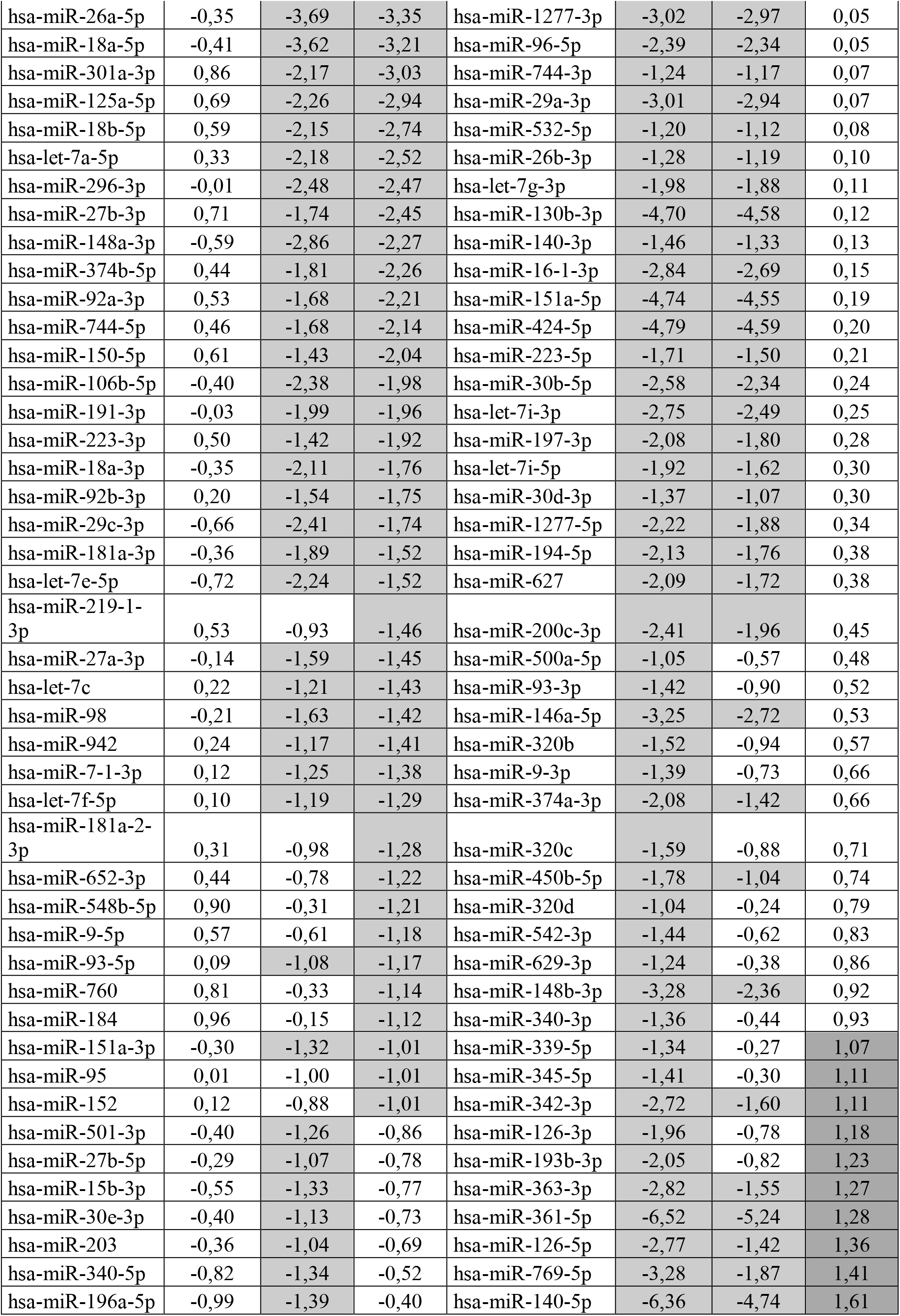
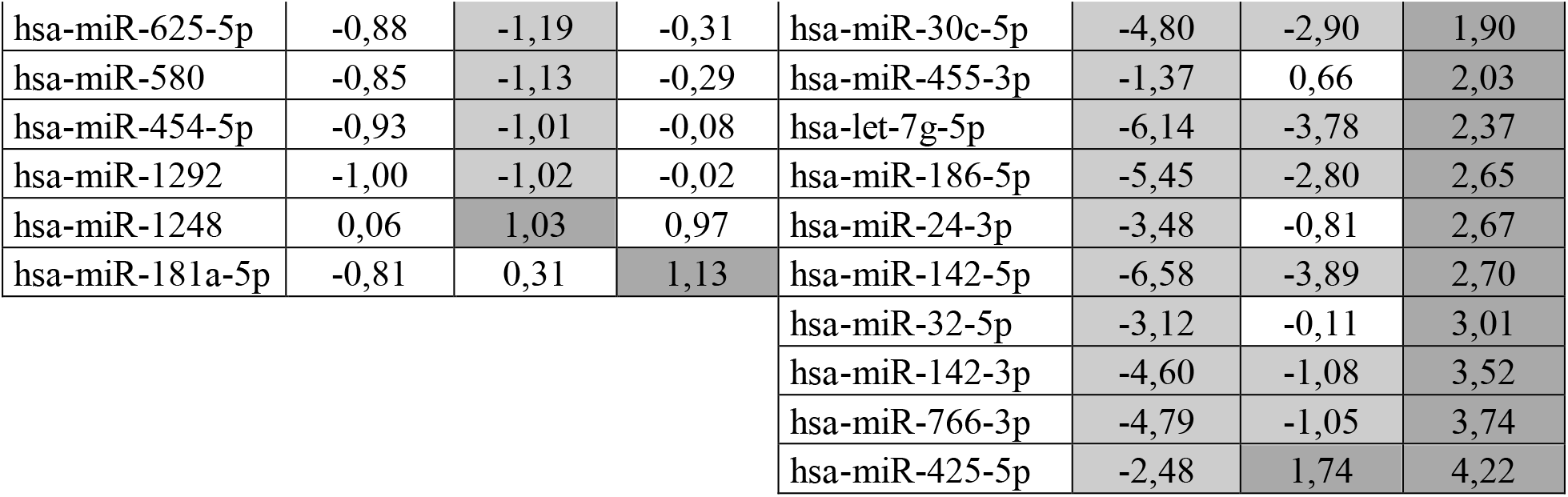
A list of miRNAs de-regulated in camptothecin-treated Jurkat cells. The cloning frequency of miRNAs was calculated as read-per-million (RPM) following the removal of the adapter sequences. The miRNAs with a cloning frequency of 10 RPM in all three samples were used in calculations. The fold of expression is presented in a log ratio. JNN, drug-free, control cells; JAN, AnnexinV-negatif fraction of drug-treated cells; JAP, AnnexinV-positive fraction of drug-treated cells. Dark and light gray represent up- and down-regulated miRNAs, respectively.

**Table 2.** A summary of differentially expressed miRNAs and qPCR validation. A. The dysreglated miRNAs from Table 1 is summarized. JNN, the contol Jurkat cells not treated with camptothecin; JAN and JAP the Annexin V-negative and positive populations of the camptothecin-treated cells, respectively. +, induced miRNAs; -, suppressed miRNAs; =, equally expressed miRNAs. B. cDNA was prepared from a fraction of the total RNAs used in deep sequencing analysis using RT^2^ miRNA cDNA kit (SA Biosciences). qPCR was performed in duplicates of three biological replicates. U6 small RNA was used for normalization. ±, standart deviation.

| JAN/JNN | JAN/JNN | JAP/JAN | # |
| --- | --- | --- | --- |
| - | - | = | 57 |
| = | - | - | 30 |
| + | = | - | 29 |
| - | - | + | 12 |
| = | - | = | 11 |
| - | - | - | 10 |
| - | = | = | 9 |
| = | = | - | 9 |
| - | = | + | 7 |
| + | - | - | 5 |
| + | + | = | 1 |
| + | = | = | 1 |
| - | + | + | 1 |
| = | + | = | 1 |
| = | = | + | 1 |

**B**
| miRNAs | Deep-sequencing |  | qPCR |  |
| --- | --- | --- | --- | --- |
|  | JAP/JNN | JAN/JNN | JAP/JNN | JAN/JNN |
| miR-7 | 11,6 | 0.7 | 0.22 ± 0.01 | 0.14 ± 0.01 |
| miR-17 | 0.07 | 0.33 | 0.27 ± 0.03 | 0.21 ± 0.03 |
| miR-18a | 0.08 | 0.75 | 0.25 ± 0.02 | 0.21 ± 0.01 |
| miR-26a | 0.08 | 0.79 | 0.21 ± 0.02 | 0.21 ± 0.02 |
| miR-93 | 0.47 | 1.07 | 0.3 ± 0.03 | 0.2 ± 0.02 |
| miR-425 | 3.35 | 0.18 | 2.06 ± 0.25 | 0.15 ± 0.01 |

## Discussion

The balance between proliferation and apoptosis is important for the overall cellular homeostatis. Apoptosis is particullarly important in modulating the fate of immune cells, including T cells. Microarray and deep-sequencing studies have been instrumental in identifying several miRNAs involved in cell proliferation or apoptosis (Subramanian and Steer, 2010). However, these studies were mainly conducted with heterogenous cancerous tissues in which the apoptotic state of the cells were not synchronized. Thus, we used the Jurkat cell line and the universal apoptosis inducer camptothecin to identify the miRNAs involved in apoptosis. The apoptotic cells were identified by marking the cells in which phosphatidylserine was exposed to the cell surface, which is readily detected by Annexin V labeling. Sorting cells based on their Annexin-V labelling allowed us to obtain the miRNA expression profile of a purely apoptotic cell population (Annexin V/7AAD-double positive).

miR-14 and bantam were the first miRNAs shown to have apoptotic function in *Drosophila* (Brennecke et al., 2003; Xu et al., 2003). Studies on various cancer cells showed a prominent p53-mediated regulation of miRNAs, particularly miR-34 family, miR-215 and 192, with pro-apoptotic capacity (He et al., 2007; Georges et al., 2008). However, we did not detect any differential expression of these miRNAs in our study. Let-7 and miR15/16 were also reported to be associated with apoptosis (Ghodgaonkar et al., 2009). miR-16-1 and let-7d, -7g and 7i were suppressed in the camptothecin resistant JAN cells. miR-16-2 and miR-15a, on the other hand, were slightly up-regulated. These miRNAs were suppressed in camptothecin-treated Jurkat cells. The miR-17-92 cluster, which encodes miR-17.3p, -17.5p, -18a, -19a, -20a, -19b, and 92, is one of the well-established miRNA clusters with antiapoptotic activity (He et al., 2007; Georges et al., 2008). We observed nearly 4-5-fold down-regulation of these miRNAs in camptotecin-treated Jurkat cells paralel to the published results. miR-125 and -128, which are reported to be anti-apoptotic miRNAs in Jurkat cells (Yamado et al., 2014; Zhou et al., 2014), are downregulated in the camptothecin-treated cells as expected. miR-143, which is expressed at an extremely low level in cancer cell lines compared to normal tissues, is involved in Fas-mediated apoptosis in Jurkat cells (Akao et al., 2009). Accordingly, this particular miRNA was nearly undetectable in Jurkat cells. However, its expression did not change in response to the camptothecin treatment.

Our study revealed the involvement of additional miRNAs that may modulate apoptosis in human Jurkat T cells (Table 1). The most striking dysregulation were detected in the expression of miR-17*, −128, −140, −142, -161, −186, −766, and −1268. Although these miRNAs are dysregulated in camptothecin-treated Jurkat T cells, we cannot unequivocally state that they are directly involved in modulating apoptotic pathways. It is possible that they may be associated with response to drug treatment such as drug efflux. Further studies are required to determine, if any, their involvement in apoptotis and related signalling pathways.

Although camptothecin is known to be a potent apoptosis inducer, a portion of the cells were unresponsive to the drug despite over 100-fold increase in the drug concentration (Fig. 1A and F). This observation was consistent with the previous reports that there may be phenotypic and genotypic heterogeneity in clonal cell populations (Stockholm et al., 2007; Walling and Shepard 2012). To validate the cellular heterogeneity in the Jurkat cell line, we sorted the Annexin-V-positive cells (JAP) from the Annexin-V-negative cells (JAN) following the initial drug treatment and re-treated the Annexin-V-negative, apoptosis-resistant cells with camptothecin. These cells were intrinsically resistant to the drug (Fig. 1F). To investigate whether the differential apoptotic response of each subpopulation is associated with the differential miRNA expression, we subjected the total RNAs from each subpopulation to deep-sequencing using the un-treated cells as control. There were 79 miRNAs commonly down-regulated in both populations (Table 2A). Quite interestingly, although some 16 and 30 miRNAs were down- and up-regulated in the drug resistant JAN sample, they were equally expressed in the drug sensitive JAP sample. More interestingly, a total of 6 miRNAs (let-7b-5p, miR-15a-5p, 324-5p, 128, 425 and 720) were reciprocally expressed in the drug-sentisive JAP and resistant JAN subpopulations (Table 1). This observation suggests that the miRNA expression data obtained from drug treatments should be carefully interpreted if the cells of the desired phenotype are not sorted from the other cells. The presence of the cells with undesired phenotype may introduce noise that may interfere with the accurate assessment of the phenotype-specific miRNA expression profiles.

## Acknowledgements

The authors wish to thank the IYTE BIOMER for the instrumental support. This research was supported by a TUBITAK grant 107T475 (to BA). The funder had no role in study design, data collection and analysis, decision to publish or preparation of the manuscript.

